# Deteriogenic flora of the Phlegraean Fields Archaeological Park: ecological analysis and management guidelines

**DOI:** 10.1101/804823

**Authors:** Riccardo Motti, Giuliano Bonanomi, Adriano Stinca

**Affiliations:** Department of Agricultural Sciences, University of Naples Federico II, Via Università 100, 80055 Portici (Naples), Italy; Department of Environmental, Biological and Pharmaceutical Sciences and Technologies, University of Campania Luigi Vanvitelli, Via Vivaldi 43, 81100 Caserta, Italy

**Keywords:** archaeological sites, biodeterioration, biological agents, bioreceptivity, conservation management, Hazard Index, higher plants, monument conservation

## Abstract

Biodeterioration, the alteration caused by living organisms, on historical buildings and stone monuments is a well-known problem affecting two-thirds of the world’s cultural heritage. The study of the flora growing on wall surface is of particular importance for the assessment of the risk of biodeterioration of stone artifacts by vascular plants, and for maintenance planning. In this study, we investigate how rock type, exposure and inclination of the wall affect the biodeteriogenic flora at 13 sites of the Archaeological Park of the Phlegraean Fields located in the province of Naples, in southern Italy. For each site, we analysed randomly selected square areas with 2 × 2 m size, representing the different vegetation types in terms of vascular plant species cover. The total number of plant species recorded was 129, belonging to 43 families. *Erigeron sumatrensis, Sonchus tenerrimus*, and *Parietaria judaica* are the most commonly reported species, while *Capparis orientalis* is the species with the highest average coverage. Substrate type, exposure and surface inclination affect the floristic composition, with the average plant cover significantly higher on vertical surfaces and at western and southern exposure. All the main biodeteriogenic vascular plant species grow on more or less porous lythotype like yellow tufa, conglomerate and bricks. Finally, woody plants eradications methods are proposed by the tree cutting and local application of herbicides, to avoid stump and root sprouting and to minimize the dispersion of chemicals in the surrounding environment.

## 1. Introduction

In recent years, increasing attention has been paid to wall flora growing on archaeological and historical sites in the Mediterranean basin (Krigas et al., 1999; Spampinato et al., 2005; Iatrous et al., 2007; Motti and Stinca, 2011; Bartoli et al., 2017; Cicinelli et al., 2018; Dahmani et al., 2018). Although plants can in some cases be considered a protective resource for monuments (Miller, 2012; Erder, et al., 2013), in most cases they pose a severe threat to their conservation (Caneva et al., 2003; Celesti-Grapow and Blasi, 2004; Tjelldén et al., 2015; Minissale et al., 2015).

Walls can be considered an extreme environment for plant life in many respects. Segal (1969) was the first to show that wall habitats show ecological features comparable with rocks in natural environments and could be described as artificial, highly selective ecosystems (Ellenberg, 1996; Laníková and Lososová, 2009; Francis, 2011).). Wall surfaces, particularly vertical sections, offer limited opportunities for root development, the accumulation of organic matter and mineral nutrients thus limiting edaphic development and, thereafter, plant establishment (Duchoslav, 2002; Francis, 2011). Physical and environmental characteristics of walls determine their capacity to act as habitat, and control the possibility of plants to colonise such man-made ecosystems. The factors which most influence the capacity of walls to function as habitat for vascular plants are wall size, construction materials, inclination, exposure and wall age (Francis, 2011).

Higher plant colonisation of stone monuments also depends on local factors such as human disturbance, microclimate in terms of temperature and humidity, and interaction with other plants (Segal, 1969; Kumbaric et al., 2012; Ceschin et al., 2016). Establishment of plant communities on walls generally depends on the level of disintegration of building materials, with the presence of crevices, fractures and interstices that promote root development and plant growth. Nevertheless, also the technology of wall building affects the growth of plant species which are able to colonise such artificial habitats (Duchoslav, 2002; Francis, 2011). Moreover, the vegetation surrounding the investigated site affects the composition and diversity of flora growing on stone structures (Duchoslav, 2002).

The Phlegraean Fields Archaeological Park (henceforth PFAF) was established in 2016 and includes 25 sites from the Graeco-Roman period spread over an area of about 8,000 hectares. The 25 archaeological sites include ancient settlements, villas, thermal baths, temples, amphitheatres and tombs. The study sites are inserted in a complex landscape with several different habitats such as coastal and lake vegetation, Mediterranean scrubland, thermophilic and mesophilic woodland, grassland and low impact farmland (Motti and Ricciardi, 2008). Therefore, the investigated sites proved to be an interesting case study due to their great floristic richness, historical value and natural context. In the present study, we investigate the role of lithotype and microclimatic factors in terms of exposure and inclination of man-made structures in controlling the occurrence and distribution of vascular plants in stone monuments.

Given the above considerations, the specific aims of the present work were to analyse the vascular flora deteriogens of the PFAF and assess the risk of structural biodeterioration. Such knowledge is essential for the purposes of preserving the cultural landscape and for choosing appropriate management practices to prevent and eradicate vascular plants so as to minimise biodeterioration.

## 2. Materials and methods

### 2.1. Study sites

The Phlegraean area includes an insular part with the islands of Procida, Vivara and Ischia and a continental area, known as the Phlegraean Fields (Fig. 1).The area as a whole presents a highly articulated geomorphological configuration. In a very small area, bounded by a long coastline with beaches and rocky headlands, numerous volcanic calderas are interspersed with small lakes and plains. The area draws its origin from the eruption of 35,000 years ago, when a huge alkali trachytic ignimbrite followed by the subsequent collapse of the ancient volcano called Archiflegreo was released (Rosi et al., 1983). This phenomenon has produced a volcanic system with a complex hilly landscape, within which each peak represents the relict of ancient volcanic edifices, craters or eruptions.

**Fig. 1.**
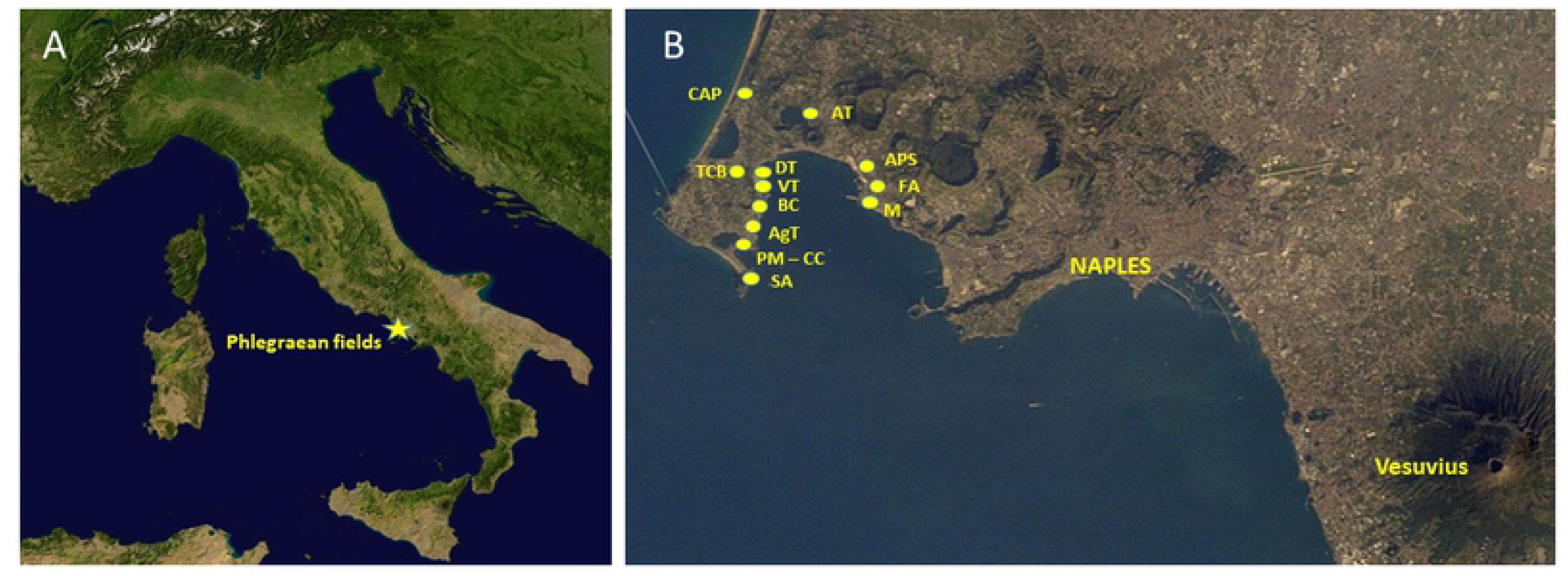
Study site (A), and location of the 13 selected sites in the PFAF (B)

Human settlements in the Phlegraean area, and especially in Cumae, date back to the III millennium BC. Founded by the Greeks in the 8th century BC, Cumae and its territory assumed great political and economic importance that allowed an expansion of its sphere of influence with the foundation of Dicearchia, the current Pozzuoli (Lombardo and Frisone, 2006). The maximum splendour of the Phlegraean area coincides with the end of the Republican age, when it became the focal point for the cultural and economic elite from Rome, and the whole territory is dotted with villas, palaces and sumptuous bath complexes (Maiuri, 1958).

The fall of the Roman Empire was followed by the decline of this area with the ruin of man-made structures already damaged by bradyseism. For many centuries, agriculture and silvi-pastoral activities were dominant, although much of the farmland and forest has been lost to extensive and chaotic urbanisation in recent decades (Motti et al., 2004).

The climate of the Phlegraean fields is influenced by both its geographical position close to the Tyrrhenian Sea and its low altitude, reaching its maximum height at Mt. Sant’Angelo alla Corbara (319 m a.s.l.). Average rainfall (863 mm) and temperature (17.0 °C) in the area are typical of a Mediterranean climate, with a hot dry period between June and August. The whole Phlegraean flora now comprises approximately 750 taxa (Motti and Ricciardi, 2005). In our study, the floristic survey concerned 13 of the 25 sites included in the PFAF (Fig. 1; Tab. 1). The remaining sites were not surveyed because they are underwater or currently inaccessible.

Tab. 1. The 13 study sites selected in the PFAF area, their abbreviations and number of surveys carried out at each site

### 2.2. Data collection and analysis

The field surveys were carried out from March to September 2018. Overall, we carried out 143 vegetation surveys (Tab. 1). The number of surveys, having taken different types of substrate into account, was proportional to the size and plant cover of each site. In each survey, we analysed randomly selected 2 x 2 m sampling units to represent the different vegetation types in terms of plant cover and floristic diversity. For each sampling units the following data were supplied: site name, position (UTM coordinates), substrate, position (vertical or horizontal), exposure and floristic list with percentage cover for each species. The plant specimens were identified in the field except for dubious cases, which were later identified at the Laboratory of Applied Ecology of the Department of Agricultural Sciences of Portici, according to Pignatti (1982), Pignatti et al. (2017a; 2017b; 2018), and Tutin et al. (1964; 1980; 1993). The nomenclature follows the checklist of Italian vascular flora (Bartolucci et al., 2018; Galasso et al., 2018). Families are organised based on APG IV (2016) for angiosperms. To evaluate the hazard of deteriogenic species, for each taxon the hazard index (HI) was assigned according to Signorini (1995, 1996). Plant life form was classified according to Raunkiaer (1934), mostly verified by field observations. The chorotype was assigned according to Pignatti et al. (2017a, 2017b, 2018). Herbarium specimens are deposited in the Herbarium Porticense (PORUN).

## 3. Results and discussion

### 3.1. Deteriogenic flora

In all, 129 plant species were recorded (Tab. S1), belonging to 43 families, of which the most species-rich are the Asteraceae (25 taxa), followed by the Poaceae (18 taxa) and Fabaceae (16 taxa).

*Erigeron sumatrensis* (HI=2) is the most commonly reported species in the 143 samples (Fig. 2), followed by *Sonchus tenerrimus* (HI=5), *Parietaria judaica* (HI=5) and *Dittrichia viscosa* subsp. *viscosa* (HI=5). Among the ten species with the maximum average cover, seven show woody habits, with a Hazard Index between 5 and 10. *Capparis orientalis* (HI=8) is the species with the highest average cover (Fig. 3) followed by *Dittrichia viscosa* subsp. *viscosa* (HI=5) and *Spartium junceum* (HI=8).

**Fig. 2.**
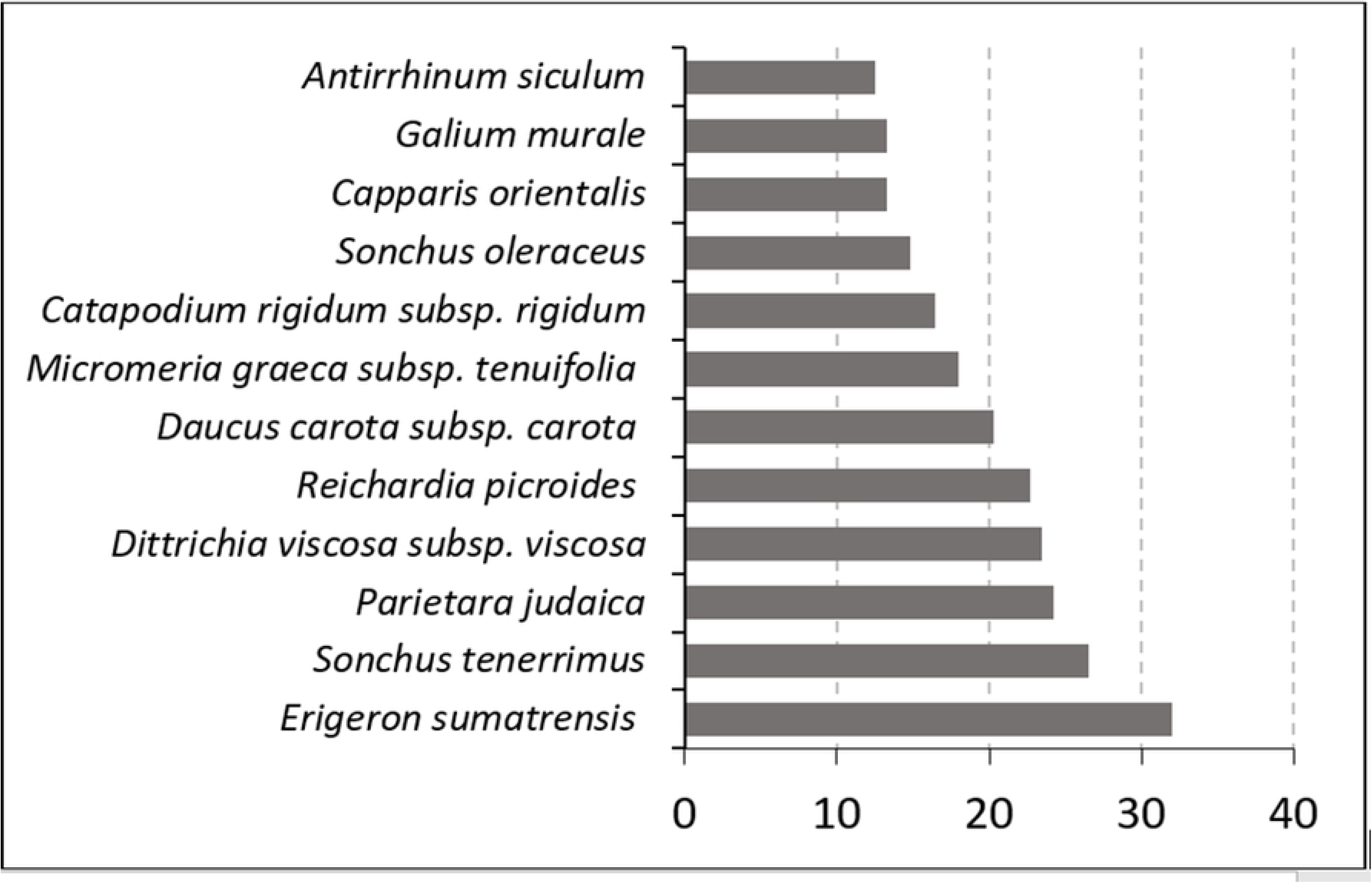
List of the 12 most commonly recorded species in the 143 sampling units (number of records for each species).

**Fig. 3.**
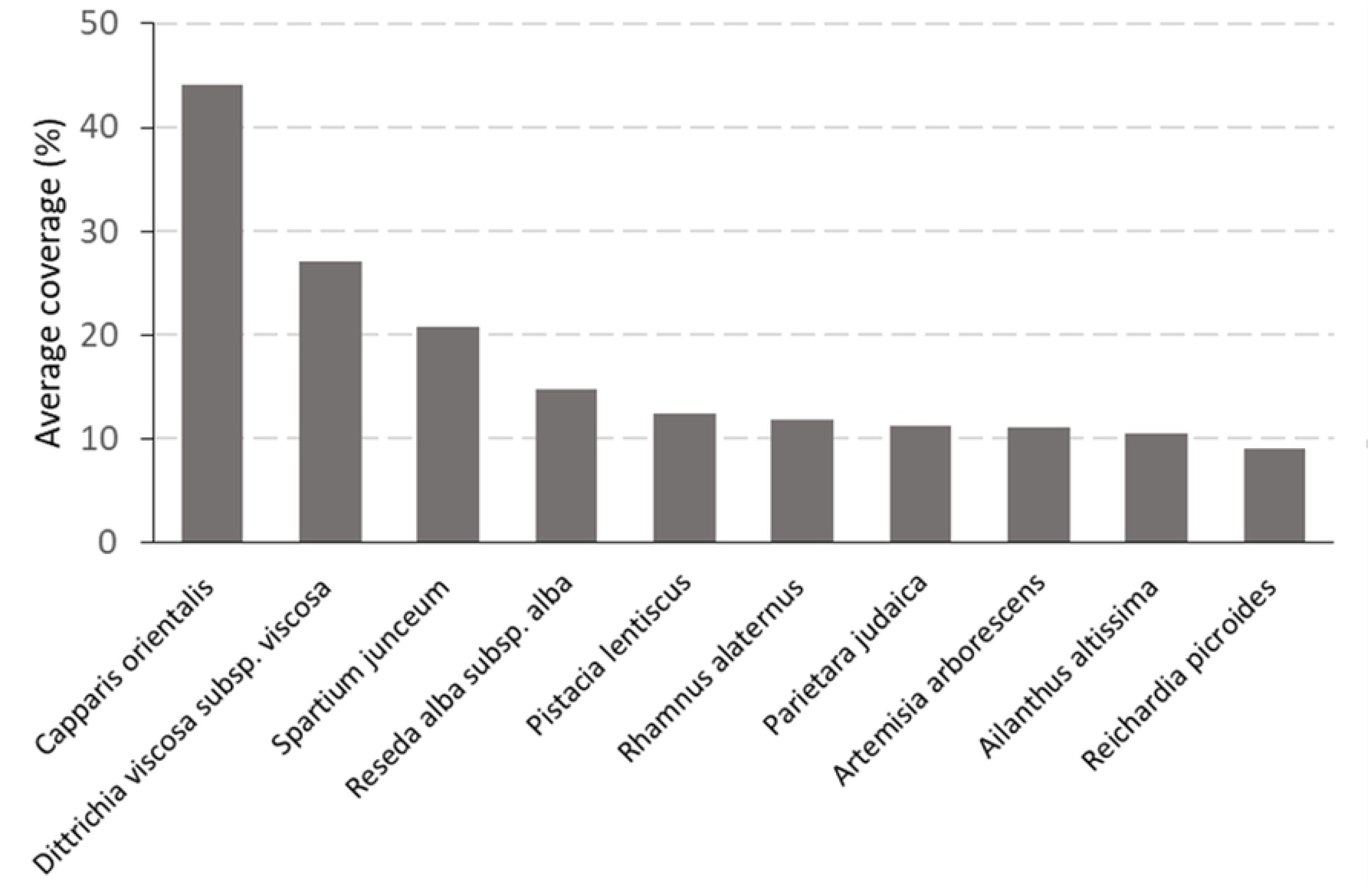
Cover of the 10 most abundant species in the 143 sampling units

The normal chorological spectrum (Fig. 4) revealed the prevalence of Mediterranean species (33.7%), of which the most representative are euri-Mediterranean (62.8%) vs. steno-Mediterranean (37.2%). Widely distributed species are well-represented (35.7%), of which alien species amount to 30.4%. These data are similar to those of the floristic list of the whole Phlegraean Fields area (Motti and Ricciardi, 2005).

**Fig. 4.**
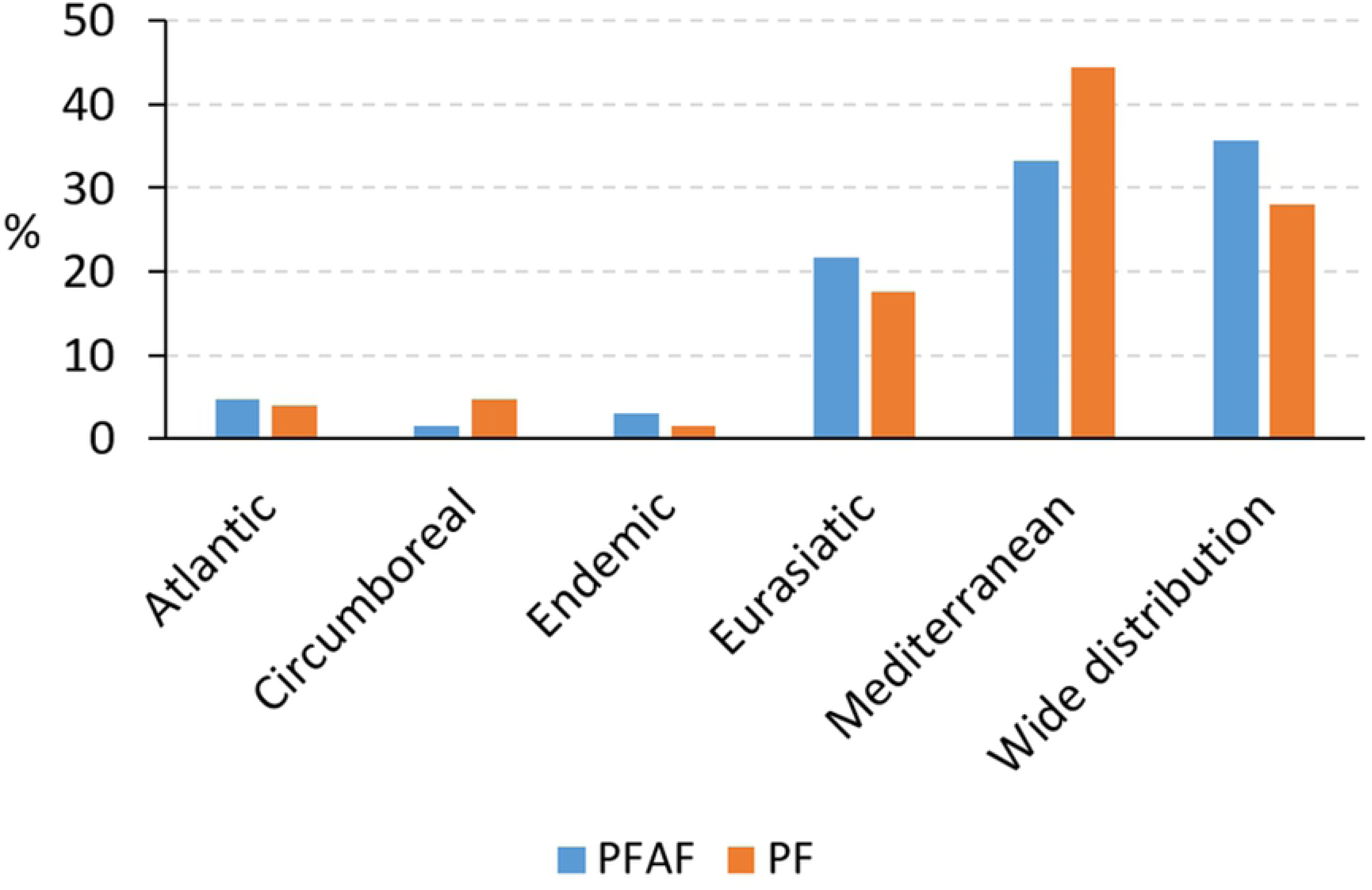
Normal chorological spectrum of the flora of the Phlegraean Fields Archaeological Park (PFAF) compared with that of the flora of the whole area of the Phlegraean Fields (PF) (Motti and Ricciardi, 2005)

The archaeological sites of the PACS are located in a floristic context dominated by species associated with agricultural environments, as well as by woody species typical of Mediterranean tufaceous coastal hill ecosystems (Motti and Ricciardi, 2008). Our data indicate that the flora growing on stone structures partially reflects this kind of vegetation. Hence the floristic composition of the PACS is influenced by its proximity to natural areas (Duchoslav, 2002; Migliozzi et al., 2010).

The life form spectrum (Fig. 5A) shows a prevalence of Therophytes (48.4%) and Hemicryptophytes (28.6%) at all study sites, but these life forms have no predominant cover (Fig. 5B). Woody life forms (Phanerophytes and Chamaephytes), which include the most deteriogenic species, account for 22.2% of the total frequency and 60.3% of total cover. The relationship between therophytes and hemicryptophytes (T/H ratio: 1.7) is influenced by human disturbance, which promotes the spread of short-lived species (Motti and Stinca 2011), as well as by climate.

**Fig. 5.**
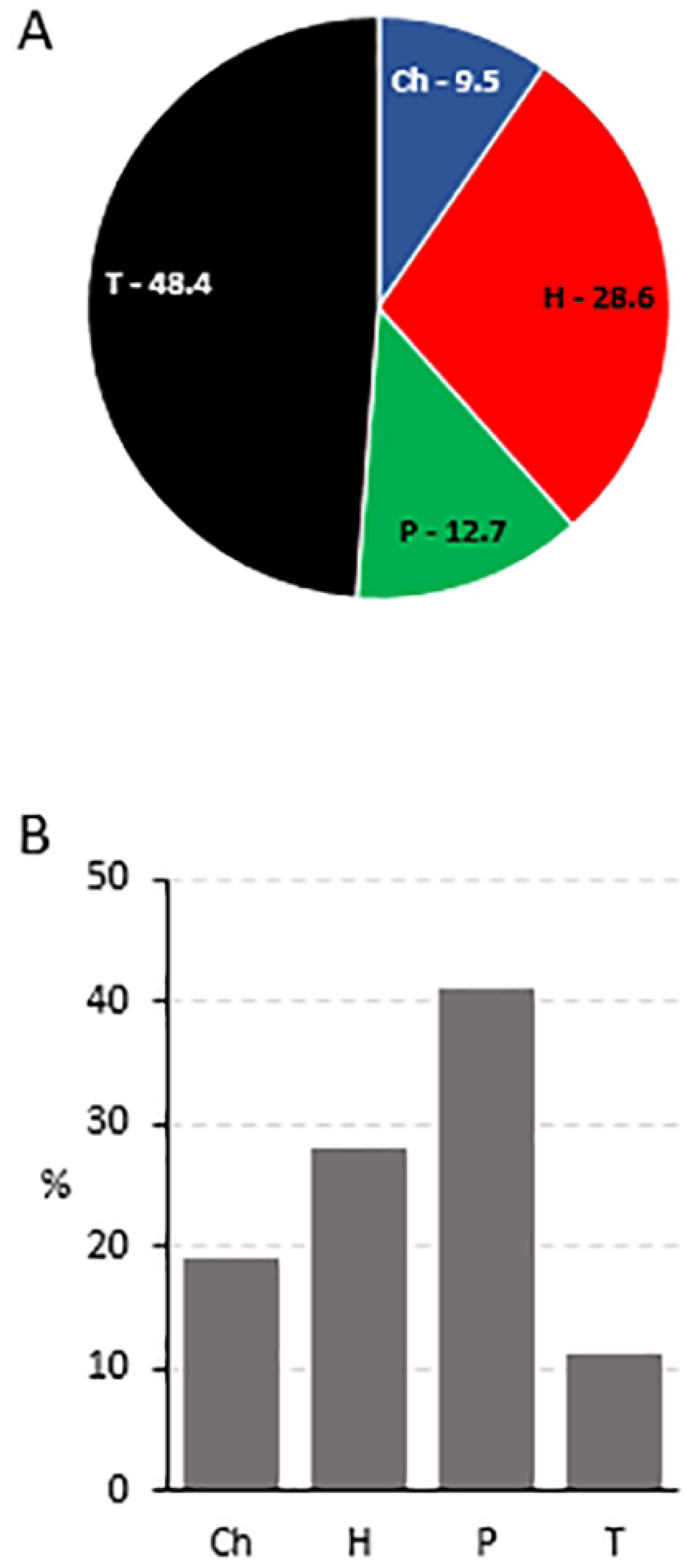
Plant life-form spectrum of the vascular flora (A) and percentage cover of different plant life forms (B) in the 143 sampling units (T=Therophytes; P=Phanerophytes; H = Hemicryptophytes; Ch = Chamaephytes).

### 3.2. Deteriogenic flora: the role of inclination, exposure and substrate type

Wall inclination affects the amount of direct solar radiation reaching the substrate and, indirectly, air and soil temperatures (Wieser and Tausz, 2007). Previous studies reported that horizontal surfaces, which provide better growing conditions, usually host a higher plant cover compared to vertical walls (Caneva et al., 1992; Lisci et al., 2003; Ceschin et al., 2016; Motti and Bonanomi, 2018). Vertical walls are often considered to be like desert habitats with a high degree of aridity, and the stone surfaces exposed to direct sunlight can reach extremely high temperatures (Garty, 1990). In contrast with previous results, in our study sites the average plant cover (Fig. 6A) was significantly higher on vertical surfaces (Fig. 7A). As shown also by Duchoslav (2002), Therophytes were more common on horizontal surfaces, while Hemicryptophytes, Chamaephytes and Phanerophytes, the most biodeteriogenic life forms, grow rather on vertical surfaces (Fig. 7B). This could be explained by the greater ability of the latter life forms to absorb water from greater depths (Kumbaric et al., 2012; Caneva et al., 2009). Alternatively, the high frequency and cover of Therophytes on flat surfaces and the widespread occurrence of woody plants on vertical surfaces could be explained by the different effort exerted for cleaning. Indeed, vertical surfaces are more difficult for workers to reach and are therefore subject to less intense and less frequent removal of vegetation.

**Fig. 6.**
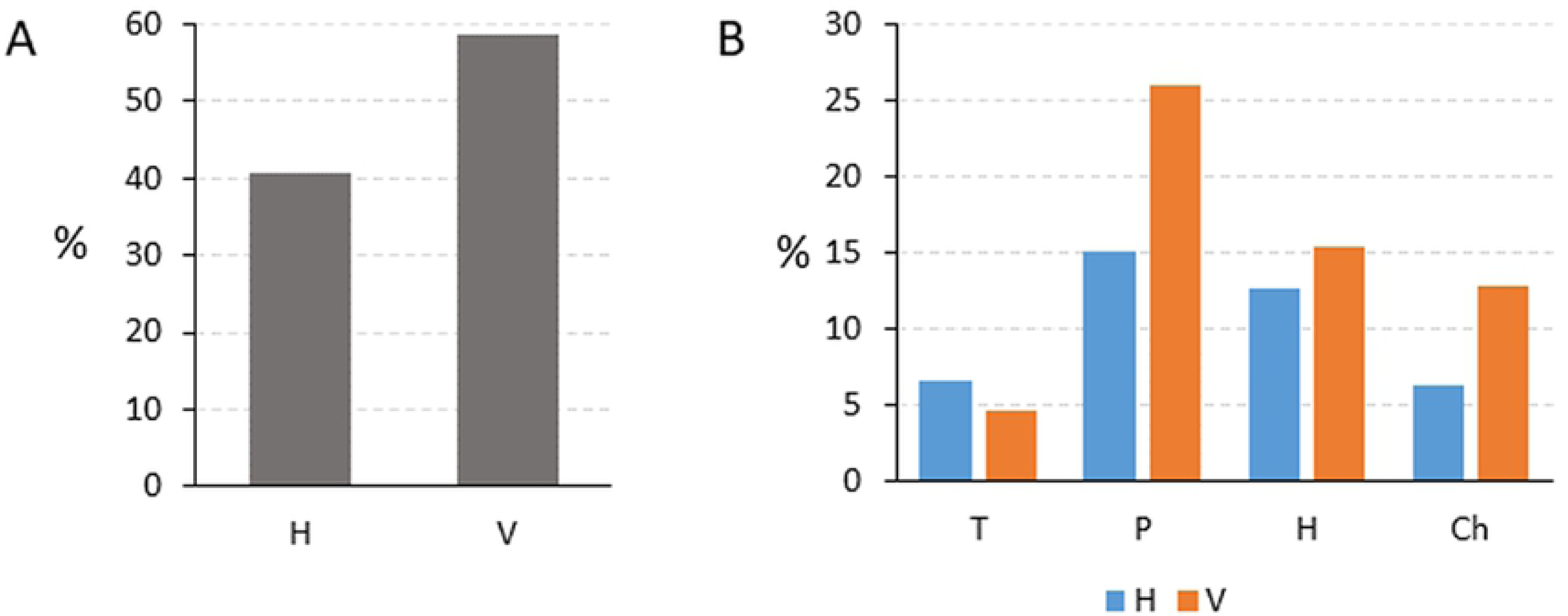
Percentage cover (A), and percentage cover of life forms (B) on horizontal (H) and vertical (V) surfaces in the study sites. Values are averages of the 143 sampling units.

**Fig. 7.**
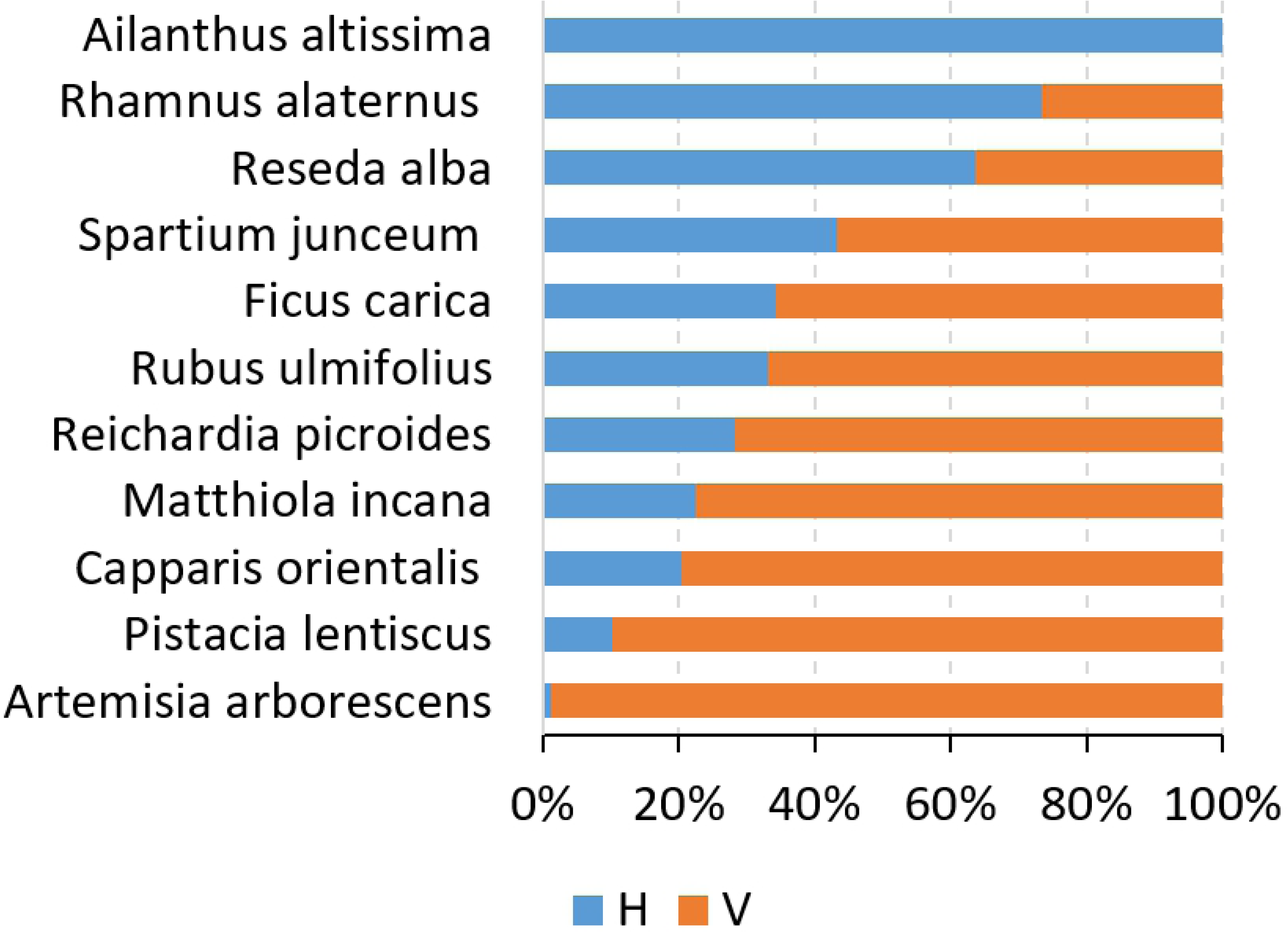
Relative cover of the 11 species with the highest HI in relation to inclination (H= horizontal; V=vertical). Values are averages in the 143 sampling units.

Among the species with the highest hazard index (HI>5), *Ficus carica, Matthiola incana, Pistacia lentiscus, Capparis orientalis, Reichardia picroides, Rubus ulmifolius* and *Artemisia arborescens* show a higher abundance on vertical surfaces. By contrast, *Spartium junceum, Rhamnus alaternus* and *Reseda alba* are almost indifferent to surface inclination, while *Ailanthus altissima* grows almost exclusively over horizontal substrates (Fig. 7).

In the Mediterranean climate context, with its long summer drought, exposure plays a major role in causing differentiation in biological colonisation (Caneva and Ceschin, 2009). At mid latitudes of the northern hemisphere, south-facing slopes receive more direct solar radiation and can be expected to be much warmer and drier than other exposures.

In the study area, plant cover was highest on western and eastern-exposed walls (Fig. 8A). Phanerophytes, which are woody plants that may reach a considerable size and have an extensive root system (Pacini and Signorini, 2009), were recorded mainly on western and eastern exposures (Fig. 8B). By contrast, herbaceous species (hemicryptophytes and therophytes) grow preferentially on south-facing slopes. The southern slopes reproduce the general life strategies of plants found in the Mediterranean climatic area where drought-avoiding annuals predominate and herbaceous perennials, which die back to the ground surface during the summer drought, are also common (Mooney and Dunn, 1970). The woody species grow under less dry exposure (East and West) and are almost all evergreen trees and shrubs, which tolerate the less intense drought that occurs over such exposures. Moreover, as highlighted by Callaway (2007), stones can act as an inanimate “nurse” structure, promoting the establishment and growth of plants and acting as a temperature buffer. In this perspective, southerly exposure may be more favourable for plant growth due to the larger amount of solar radiation, especially during autumn and winter (Motti and Bonanomi, 2018).

**Fig. 8.**
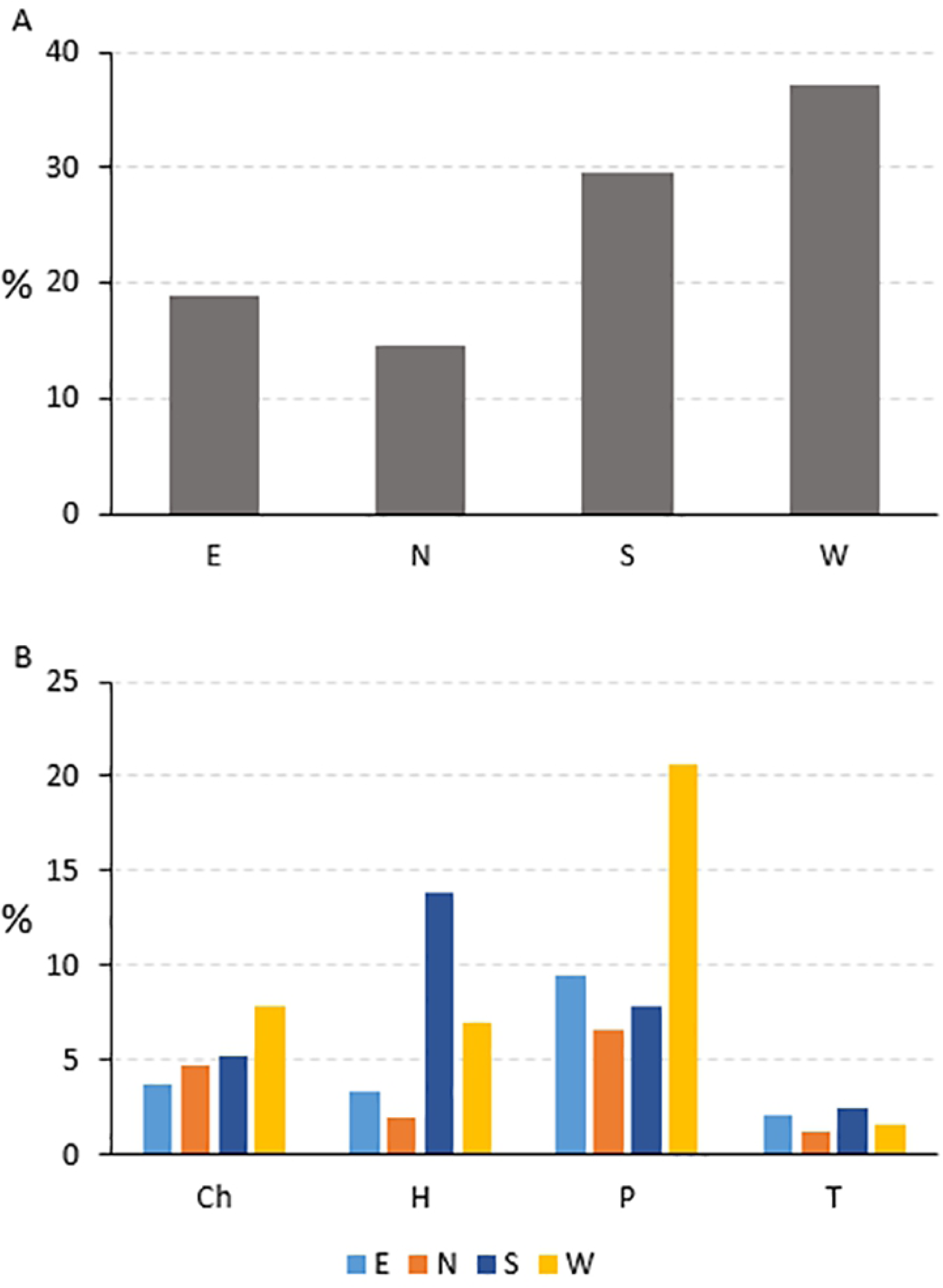
Plant cover (A, average of the 143 sampling units) and cover of different life forms (B) in relation to exposure

Natural stones are the main element of the archaeological heritage and are subject to biodeterioration: they are mostly located outdoors and the processes of deterioration affecting them are the same that play an essential role in pedogenesis (Pinna and Salvadori, 2009). Effusive magmatic rocks such as yellow tuff, piperno and basalt represent the most common stony substrates in the PFAF. Instead, the man-made structures mainly consist of the following materials: i) *opus reticulatum*, consisting of a sand and lime mortar mix into which diamond-shaped bricks of tuff were positioned (Wilson, 2006); ii) *opus latericium* walls, built with clay-fired bricks bonded with mortar. In both cases, the bricks constitute the external parts of the wall, while the inner section is filled with a conglomerate of mortar, tuff and lapillus (Talamo P., *in verbis*).

Other non-effusive stones can be found in the study area, including marble and vitreous mosaics. Since tuff and mortar have a relatively high porosity (Kumbaric et al., 2012), they allow higher water penetration and are more likely to retain moisture compared to other lithotypes. On the above basis, we could partially explain the differences in plant cover among different substrates (Fig. 10A), which reaches the highest values on yellow tuff followed by conglomerate, *opus latericium* and *opus reticulatum*.

**Fig. 9.**
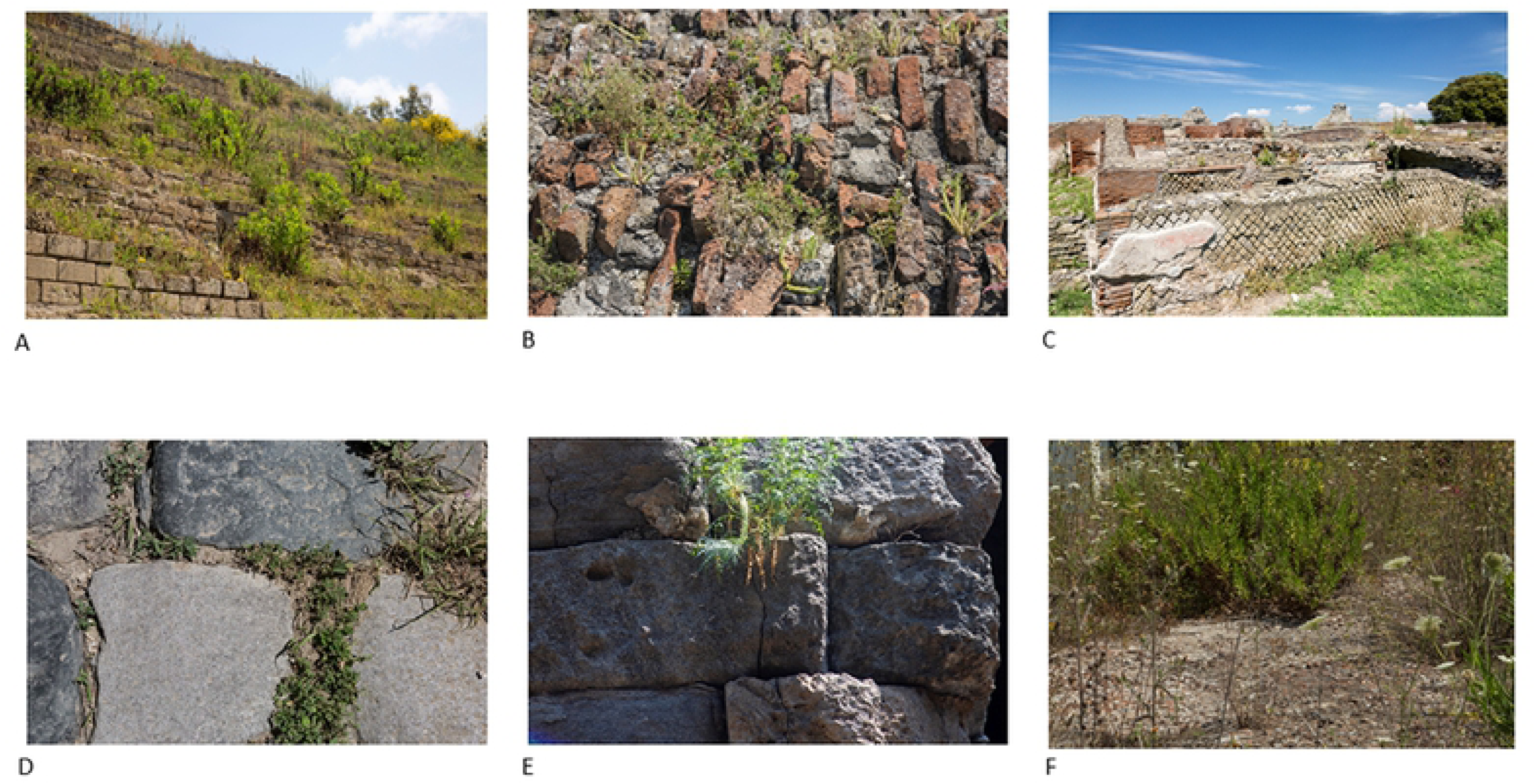
Selected images of the most common substrates found in the PFAF (A=Yellow tuff; B= *Opus latericium*; C=*Opus reticulatum*; D=Basalt; E=Piperno; F=Conglomerate).

**Fig. 10.**
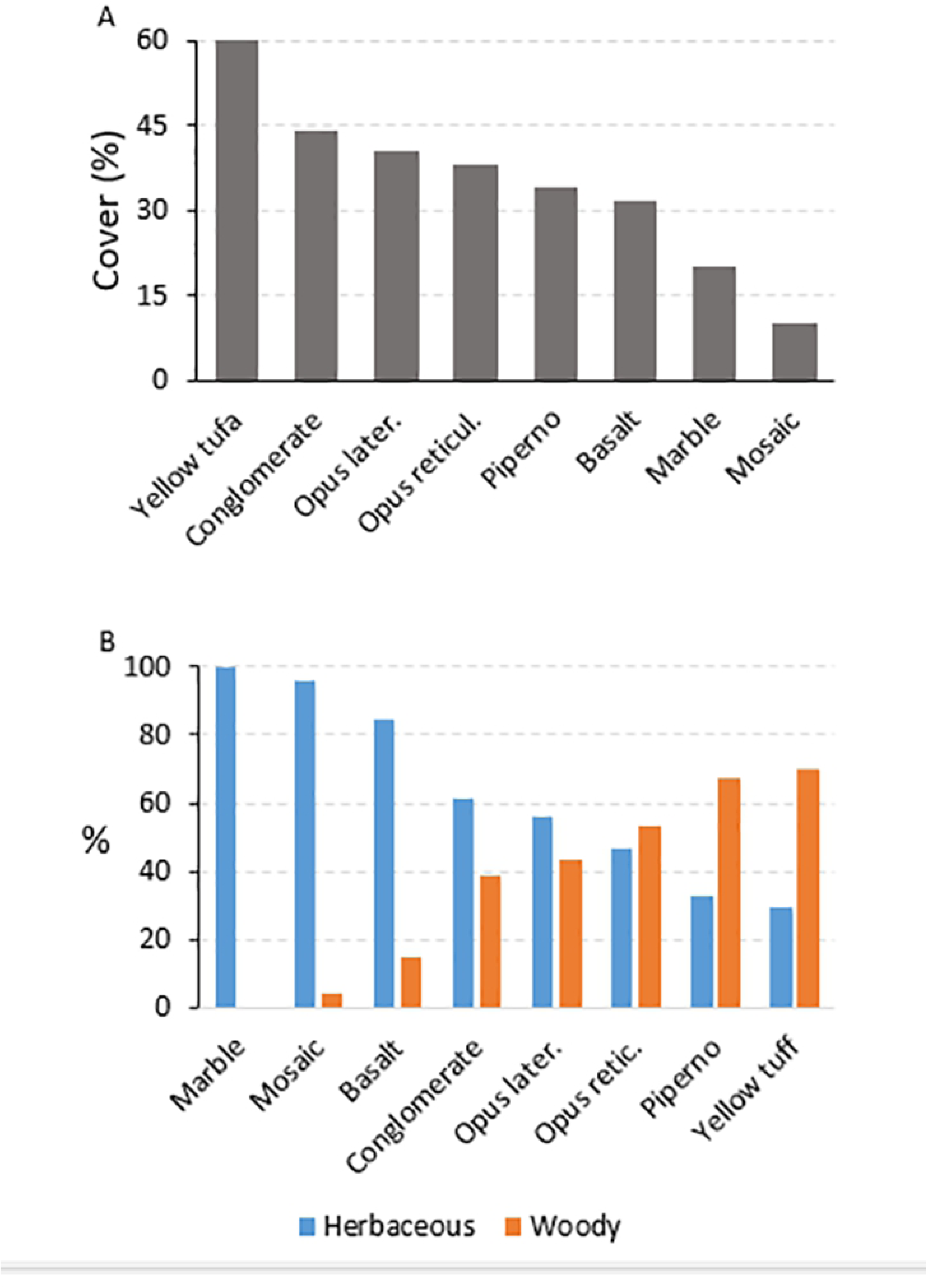
Plant cover (A) and growth habit cover (B) in relation to substrate. Values are averages from the 143 sampling units

As shown in Fig. 9B, the less porous lithotypes like marble, basalt and mosaic are mainly colonised by herbaceous species, while woody species grow preferentially on volcanic rocks and structures with a higher moisture content. We speculate that therophytes can adapt to hard and non-porous substrates because their vegetative period is limited to autumn and winter when water shortage is not a limiting factor. Instead, perennial plants require porous substrates that store water, thus allowing survival also during the summer drought.

All the main deteriogenic vascular plant species grow on more or less porous substrates (Fig. 10): none of them thrive on marble, basalt or mosaics. Some species (e.g. *Rubus ulmifolius, Rhamnus alaternus*) are quite indifferent to the lithotype, while others, such as *Ailanthus altissima, Spartium junceum, Matthiola incana* and *Artemisia arborescens*, preferentially grow on yellow tuff.

Plant growth on walls can be therefore interpreted as a dynamic process with weeds that may alter the physical conditions of the substrate in which they thrive (Fisher 1972), also through progressive disintegration of building materials. Biochemical deterioration results from assimilatory processes, where the organism uses the stone surface as a source of nutrition, and from dissimilatory processes, where the organism produces a variety of metabolites that react chemically with the stone surface (Mortland et al., 1956; Caneva and Altieri, 1988). Carbon dioxide, produced through roots respiration, changes into carbonic acid [H_2_CO_3_] in an aqueous environment. The carbonic acid reacts with calcium carbonate [CaCO_3_] and magnesium [MgCO_3_] insoluble present in several substrates, forming calcium bicarbonate [Ca(HCO_3_)_2_] and magnesium [Mg(HCO_3_)_2_] soluble (Mishra et al., 1995; Pinna and Salvadori, 2005).

Plants exploit and help to create microenvironments suitable for plant growth, (Allsopp et al., 2004).and pre-existing plant cover favours the establishment of other taxa, also protecting them against evaporation and regulating relative humidity (Segal, 1969). In this sense, the first plants that colonise the walls mainly have a herbaceous growth habit and could be considered pioneer species playing a key role in stone weathering: their strong fasciculate root system creates or widens crevices in which soil is formed, providing organic matter and nutrients that promote succession of typical vegetation for the biogeographic region concerned (Segal, 1969; Duchoslav, 2002).

### 3.3. Plant deterioration and management guidelines

The plant average cover for each site is shown in Fig. 12, from which it can be seen that sites like the Temple of Apollo, Flavian Amphitheatre, Baia Castle, the Sacellum and Piscina Mirabilis need urgent maintenance to eliminate the most deteriogen vascular plants so as to minimise the risk of severe structural damage. In the other sites of the complex, although the plant cover is lower, periodic assessment of case-by-case situations is required.

**Fig. 11.**
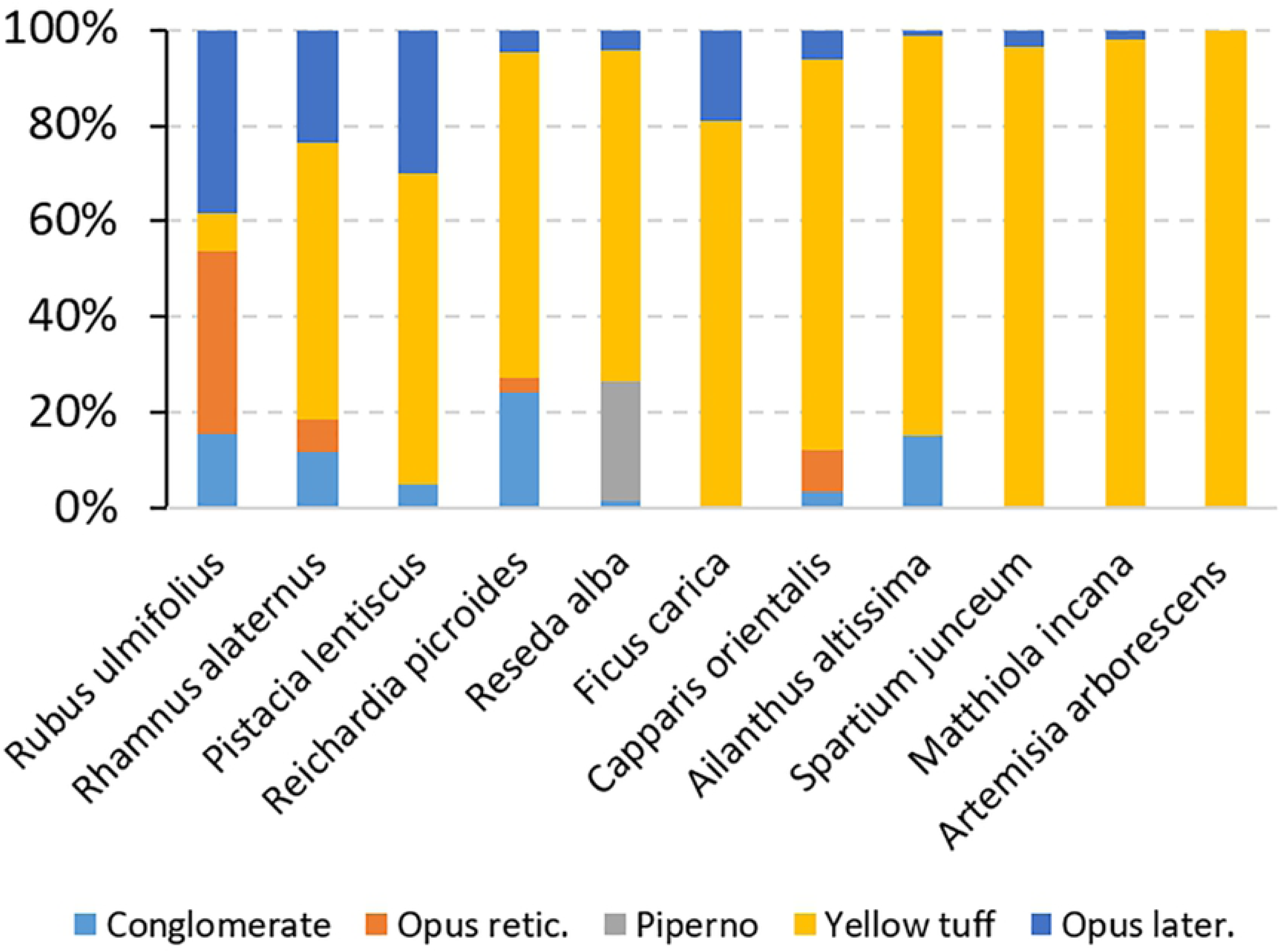
Relative cover of the 11 species with the highest HI in relation to substrates. Values are averages of the 143 sampling units.

**Fig. 12.**
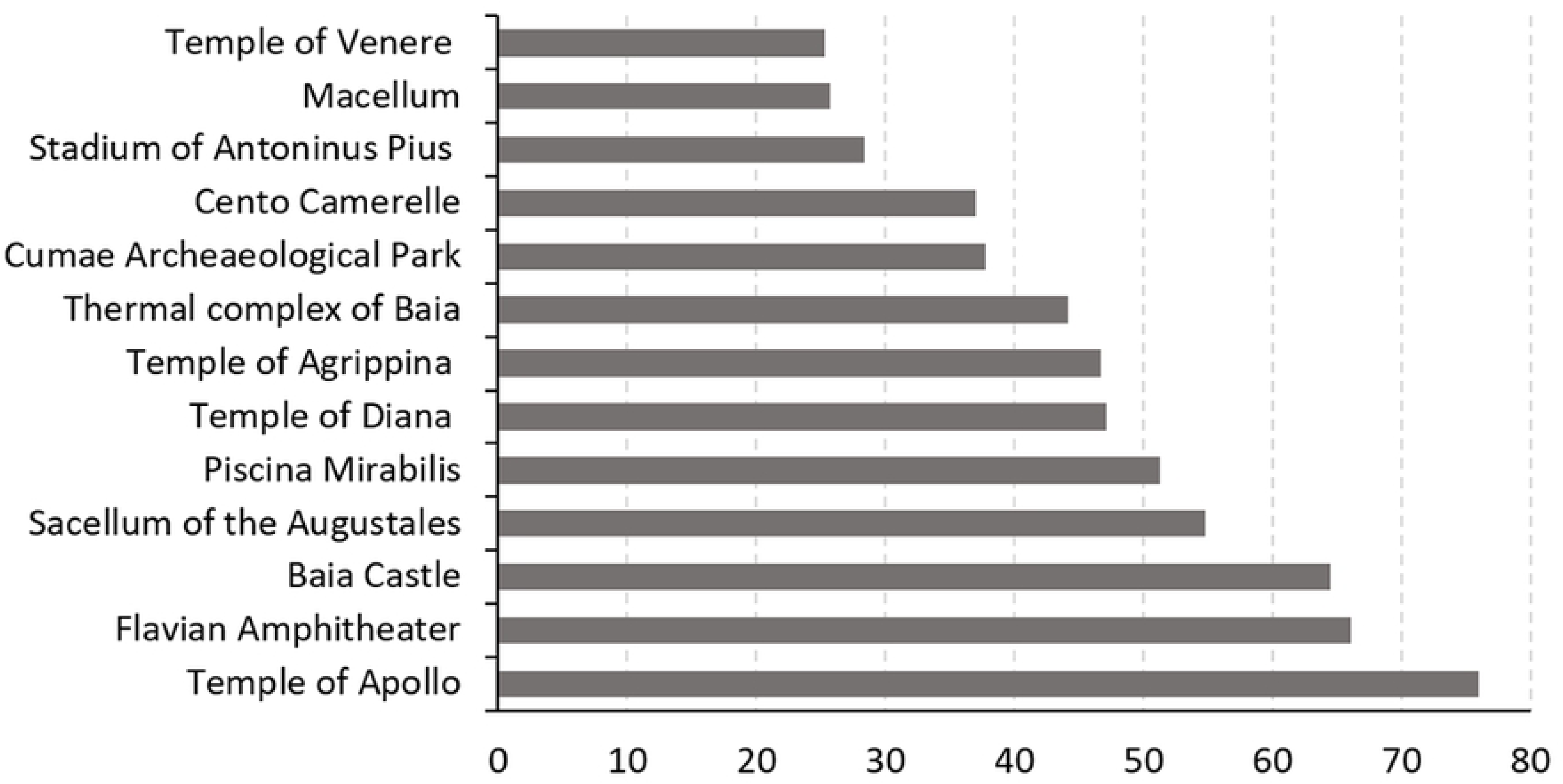
Plant cover (%) in the 13 study sites

Weed control in archaeological sites is complex and costly since proper conservation of man-made structures, the environment and the natural landscape has to be taken into account. Current practice at PFAF sites to manage undesirable vegetation relies mainly on mowing by the use of brush-cutters. This practice is strongly discouraged by archaeologists as it can cause additional deterioration to walls. Moreover, many species, especially shrubs and trees, are not completely eliminated because only the above-ground portion is cut. In recent years alternative non-chemical methods for weed control, including flame weeding and soil solarization, have been proposed (Papafotiou et al., 2016; Papafotiou et al., 2010). These treatments are often more expensive than chemical weed control due to higher treatment frequency and greater energy consumption than chemical weed control, resulting in a lower cost/benefit ratio (Kempenaar et al., 2002). Selective use of herbicides is, in our opinion, the most efficient and least costly practice for controlling and eradicating woody vascular plants, especially on vertical surfaces.

In many woody plants (e.g. *Ailanthus altissima, Capparis orientalis*) manual cutting generally stimulates stump and root sprouting due to the loss of apical dominance, such that cutting must be followed by local herbicide treatment (Caneva et al., 1996; Burch and Zedaker, 2003). The techniques suggested for this purpose are tree cutting and local application of herbicide by injection or by painting, allowing the herbicide to translocate throughout the roots and/or rhizome of the plant (Mendes et al., 2017) while maintaining the integrity of the remaining plant community.

These techniques involve the application of chemicals (e.g. glyphosate, imazapyr) directly on the plant, with no dispersion in the surrounding environment and minimising product quantities. Stem injection consists in making a cut (or a hole by drilling) downward at an angle of ∼45 degrees through the bark, 4 to 8 cm long, and then injecting a small amount of herbicide (DiTomaso and Kyser, 2007). The cut stump method, instead, involves cutting off the plant completely at its base using a chainsaw. The herbicide solution is then painted onto the exposed surface.

The choice of the most appropriate technique is made on the basis of the structural and physiological features of the species, age and size of the specimen, and the position of the plant in relation to the wall. According to Caneva et al. (2009), these practices should be followed by wall consolidation because, with the death of the living roots, collapses and structural damage could arise.

## 4. Conclusions

Our data provided useful information for understanding the role of abiotic factors (substrate, position, exposure) in determining plant growth. The eradication methods proposed in the present paper constitute an example of a multidisciplinary approach to restoration practices, in which collaboration between agronomists, archaeologists and masonry experts is desirable. The ultimate goal is to apply efficient techniques with no undesirable side effects on the substrate.

## Acknowledgements

We are very grateful to Paolo Giulierini, Director of the PFAF, and the technical-administrative staff who shared knowledge with us and for their logistical support. The authors wish to thank Pierfrancesco Talamo, archaeologist at the PFAF, for information concerning man-made structures. Stefano Emrick collaborated in field research. Mark Walters revised the English language version.

